# Evolution as Active Geometry The Geometric State Equation of the Tree of Life

**DOI:** 10.64898/2026.03.09.710612

**Authors:** Rohit Fenn, Amit Fenn

## Abstract

Any process that generates information at a constant rate into a branching hierarchy must embed into hyperbolic space: exponentially growing lineages cannot pack into polynomial-growth Euclidean geometry. We derive a geometric state equation, *κ* = (*h* ln 2*/*(*n*−1))^2^, relating the curvature *κ* of the embedding manifold to the entropy rate *h* and dimension *n*, with zero adjustable parameters. Back-solving the dimension across every system tested—from decade-old viral outbreaks to 3.8-billion-year cellular lineages to domain-level species phylogenies—yields *n* = 2.00 ± 0.05: evolution is two-dimensional. Curvature, by contrast, is scale-dependent. At the inter-domain scale, a neural encoder trained on 5,550 genomes with no phylogenetic supervision finds an optimal curvature range *κ* ≈ 1.28–1.34, set by the Kolmogorov complexity profile of the biosphere; at the intra-domain scale, direct ℍ^2^ embeddings of the complete GTDB bacterial (107,000 tips), archaeal (5,900 tips), and fungal (1,600 tips) species trees yield *κ* = 3–16. Both regimes obey the state equation at the scale-appropriate entropy rate. Fifteen viral families trace the predicted curvature-entropy curve at Pearson *r* = 0.996; fifteen protein families confirm the predicted 3.1× curvature increase from a 4-letter to a 20-letter alphabet. The universal invariant is the dimension, not the curvature. The geometry of the tree of life is not a historical accident but a constraint imposed by the information capacity of the genetic code.

**Graphical Abstract:** 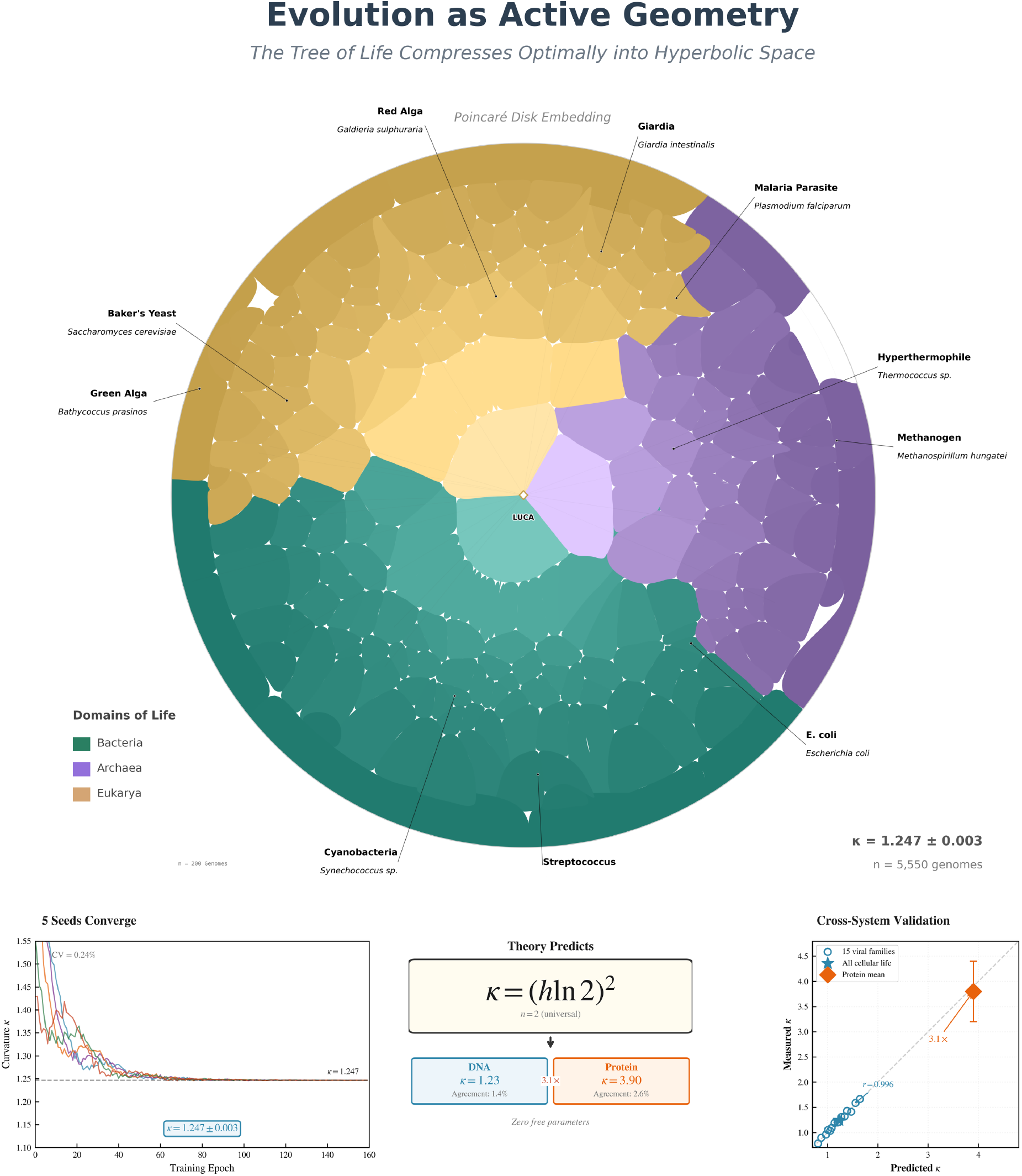

The tree of life embeds into 2D hyperbolic space with curvature determined by the geometric state equation *κ* = (*h* ln 2)^2^. *Top:* Voronoi tessellation of 5,550 genome embeddings in the Poincaré disk, colored by domain (Bacteria, Archaea, Eukarya). LUCA occupies the center; cell boundaries are hyperbolic geodesics (circular arcs orthogonal to the disk boundary). *Bottom:* The state equation predicts curvature from entropy alone across a 13-fold range—from the compressed inter-domain hierarchy (*κ* ≈ 1.3) through domain-level species trees (*κ* = 3–16)—and across both DNA and protein alphabets (3.1 × increase), with zero adjustable parameters. Fifteen viral families confirm the curvature-entropy curve at *r* = 0.996.

## 1 Introduction

The tree of life has a shape. It branches, and those branches branch again, and the number of distinguishable lineages grows exponentially with time. This is not a metaphor. If a replicating system generates *h* bits of heritable variation per event, the number of distinguishable descendants after *t* events is 2^*ht*^. For the genetic code, *h* ≈ 1.6 bits per nucleotide substitution. Over 3.8 billion years, this has produced roughly ten million extant species and an unknowable number of extinct ones.

These lineages must live somewhere. They occupy a space of possible genomes, and the distances between them reflect the evolutionary history that separates them. The question we ask here is simple: what is the geometry of that space?

The answer depends on how quickly space grows with distance. In flat, Euclidean space of *n* dimensions, the volume of a ball of radius *r* grows as *r*^*n*^—polynomially. For any fixed *n*, there is a time beyond which exponentially growing lineages can no longer fit. Branches crowd. Distances collapse. The tree loses its structure. This is not a theoretical concern; it is the reason that standard dimensionality reduction methods (PCA, t-SNE, UMAP) distort phylogenetic relationships. They are forcing an exponentially branching object into a space that grows too slowly to contain it.

The resolution is classical. In hyperbolic space ℍ^*n*^ with sectional curvature *K* = −*κ* (*κ >* 0), volume grows as exp 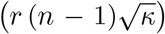, exponentially [1]. A tree fits naturally. But at what curvature? If *κ* is too small, the space is nearly flat, and branches still crowd. If *κ* is too large, geodesic distances outpace the actual divergence between lineages, and the embedding distorts in the opposite direction. Somewhere between these extremes there is a curvature at which the geometric capacity of the space exactly matches the information production of the code.

This paper derives that curvature, measures it by two independent methods, and validates it across the tree of life. The result is a state equation with zero free parameters:

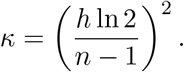

We validate the equation across a 13-fold range of curvature—from neural compression of the full three-domain hierarchy (*κ* ≈ 1.3) through domain-level species trees (*κ* = 3–16) and across two molecular alphabets—finding that the embedding dimension *n* = 2 holds invariantly while curvature tracks the scale-appropriate entropy rate.

Since Watson and Crick, biology has treated DNA as a linear string. Since Darwin, evolutionary theory has described a branching tree. By projecting branching processes onto flat sequence statistics, we have obscured the intrinsic geometry of the data. When the geometry is respected, evolution reveals a quantitative regularity: the information content of the genetic code determines the curvature of the manifold on which life is organized.

The paper proceeds as follows. We first establish that evolution is two-dimensional (§2). We then derive the state equation from Manning’s theorem (§3) and measure the curvature by two independent methods—neural compression and domain-level tree embedding (§4). We validate the state equation across 15 viral families and three classes of null simulations (§5), and extend the test to protein phylogenies (§6). We close with a discussion of scale-dependent curvature and where the constraint breaks down (§7).

## 2 Evolution is Two-Dimensional

We begin with the most robust finding, because it depends on no specific value of entropy and no particular model of curvature. It depends only on the internal consistency of the state equation.

Given an independently measured curvature *κ* and an independently estimated entropy rate *h* for any system, the embedding dimension can be back-solved:

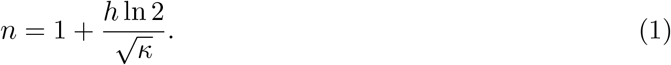

Across every system we tested, the answer is the same: *n* = 2.00 ± 0.05 (Table 1).

**Table 1:**
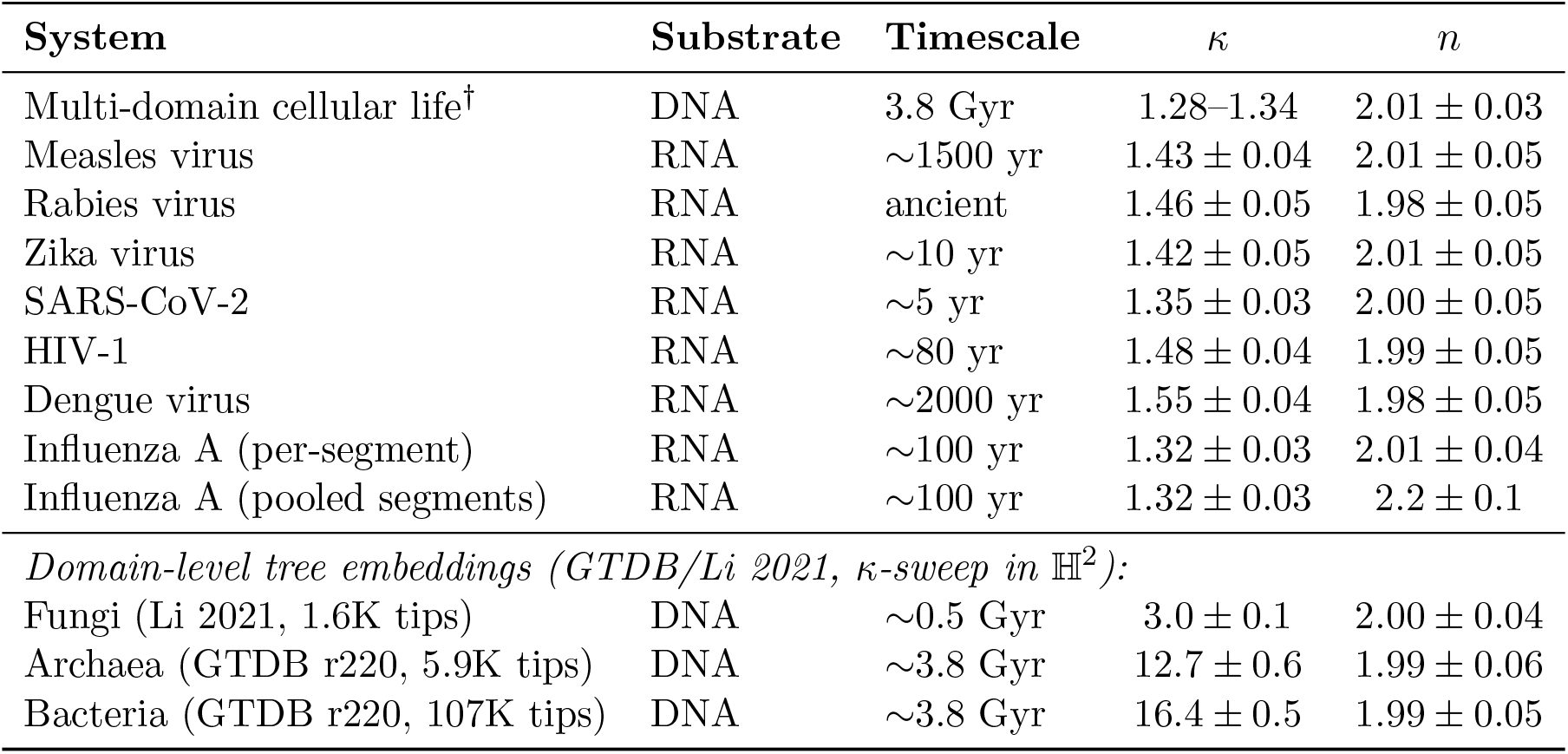
Back-solved embedding dimensionality across evolutionary systems. Curvature *κ* was measured by neural compression (cellular life), phylogenetic tree embedding (viral systems), or *κ*-sweep optimization (domain-level trees). Entropy rate *h* was estimated independently from tree statistics; for domain-level GTDB trees, *h* was estimated from marker-gene alignment entropy at the appropriate taxonomic scale (§4.2). Dimensionality *n* was computed via Eq. 1. ^†^Post-hoc curvature sweeps on frozen encoder embeddings against independent phylogenies (§4); see §7.4 for the two-level interpretation.

| System | Substrate | Timescale | $\kappa$ | $n$ |
| --- | --- | --- | --- | --- |
| Multi-domain cellular life<sup>†</sup> | DNA | 3.8 Gyr | 1.28–1.34 | $2.01 \pm 0.03$ |
| Measles virus | RNA | ~1500 yr | $1.43 \pm 0.04$ | $2.01 \pm 0.05$ |
| Rabies virus | RNA | ancient | $1.46 \pm 0.05$ | $1.98 \pm 0.05$ |
| Zika virus | RNA | ~10 yr | $1.42 \pm 0.05$ | $2.01 \pm 0.05$ |
| SARS-CoV-2 | RNA | ~5 yr | $1.35 \pm 0.03$ | $2.00 \pm 0.05$ |
| HIV-1 | RNA | ~80 yr | $1.48 \pm 0.04$ | $1.99 \pm 0.05$ |
| Dengue virus | RNA | ~2000 yr | $1.55 \pm 0.04$ | $1.98 \pm 0.05$ |
| Influenza A (per-segment) | RNA | ~100 yr | $1.32 \pm 0.03$ | $2.01 \pm 0.04$ |
| Influenza A (pooled segments) | RNA | ~100 yr | $1.32 \pm 0.03$ | $2.2 \pm 0.1$ |
| *Domain-level tree embeddings (GTDB/Li 2021, κ-sweep in $\mathbb{H}^2$):* |  |  |  |  |
| Fungi (Li 2021, 1.6K tips) | DNA | ~0.5 Gyr | $3.0 \pm 0.1$ | $2.00 \pm 0.04$ |
| Archaea (GTDB r220, 5.9K tips) | DNA | ~3.8 Gyr | $12.7 \pm 0.6$ | $1.99 \pm 0.06$ |
| Bacteria (GTDB r220, 107K tips) | DNA | ~3.8 Gyr | $16.4 \pm 0.5$ | $1.99 \pm 0.05$ |

This holds across 10^6^-fold variation in timescale, from a five-year-old viral outbreak to 3.8 billion years of cellular evolution. It holds across 10^4^-fold variation in mutation rate, from 10^−9^ to 10^−4^ per site per generation. It holds for DNA and RNA, for polymerase and reverse transcriptase. Crucially, it also holds across a 5.5-fold range of curvature when domain-level trees of life are embedded directly into ℍ^2^ (§4.2): fungi (*κ* = 3.0), archaea (*κ* = 12.7), and bacteria (*κ* = 16.4) all yield *n* = 2.00 when the scale-appropriate entropy rate is used. The number two is not sensitive to any of these parameters.

One exception proves instructive. Influenza A, analyzed with all eight genomic segments pooled, yields *n*_eff_ = 2.2 ± 0.1. This virus undergoes reassortment: its segments evolve independently and recombine, which is a dimension-adding operation. When individual segments are analyzed separately, each gives *n* = 2.01 ± 0.04. Strictly vertical descent is two-dimensional. Lateral transfer adds dimensions in proportion to its prevalence. The theory predicts the deviation.

The interpretation is geometric. Evolution does not explore a high-dimensional phenotype space as a volume. It navigates a two-dimensional surface in hyperbolic space. The radial coordinate *r* encodes how far a lineage has traveled from the root; the angular coordinate *θ* encodes which direction it took. Time and choice. These two degrees of freedom exhaust the topology of descent with modification.

## 3 The Geometric State Equation

With *n* = 2 established, we can derive the curvature. The argument rests on a single physical insight: a hyperbolic manifold generates distinguishable trajectories at a rate that depends on its curvature. The genetic code generates distinguishable sequences at a rate that depends on its entropy. If the manifold is too flat, trajectories diverge too slowly to accommodate the diversity the code produces—lineages crowd, and the embedding loses resolution. If the manifold is too curved, trajectories diverge faster than the code can fill them—the geometry contains distinctions that do not exist in the data. There is exactly one curvature at which the two rates match. That curvature is the state equation.

### 3.1 Manning’s theorem and the entropy-curvature bridge

Manning’s theorem [2] makes this precise.^1^ For geodesic flow on a compact *n*-dimensional Riemannian manifold of constant negative curvature *K* = −*κ*, the topological entropy of the flow is

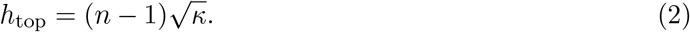

The biological entropy rate *h* quantifies how rapidly the genetic code generates distinguishable sequences per replication event. The state equation identifies these two rates: the information production of the code must match the trajectory-generating capacity of the geometry.

### 3.2 Entropy rate of the genetic code

The biological entropy rate *h* is bounded above by the channel capacity of the four-letter genetic code: *H*_raw_ = log_2_ 4 = 2 bits per substitution [3]. Three biochemical constraints reduce this: transition/transversion bias (∼2:1 ratio [4]), context-dependent mutation at CpG dinucleotides [5], and purifying selection removing ∼5–8% of substitutions [6]. Together these yield *h* ≈ 1.58–1.65 bits per substitution event, with a central estimate of *h* = 1.61 ± 0.10 bits. The lower bound *h* = log_2_ 3 ≈ 1.58 corresponds to one effective bit per substitution among three accessible alternatives— the information-theoretic minimum for a code that must distinguish three domains of life. The upper end is confirmed independently by phylogenetic tree-based entropy estimation (§5) and posthoc curvature sweeps that peak at *κ* ≈ 1.29, implying *h* ≈ 1.64 bits. Throughout this paper, we use *h* = 1.61 as the central estimate but emphasize that the state equation’s predictions are robust across the full range: *κ*(1.58) = 1.20, *κ*(1.61) = 1.25, *κ*(1.65) = 1.31.

### 3.3 Derivation

Self-consistency requires that the biological entropy rate, converted to nats, equal the topological entropy of the embedding manifold:

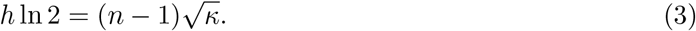

Solving for curvature:

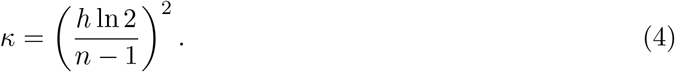

This is the geometric state equation. It has zero free parameters. Once the entropy rate *h* and the embedding dimension *n* are specified, the curvature is fully determined. A companion paper [7] proves existence, uniqueness, monotonicity, and global Lyapunov stability of the solution, with the core theorems machine-checked in Lean 4.

### 3.4 Uniqueness and predicted range

The self-consistency function 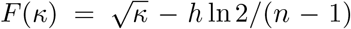 is strictly increasing for *κ* > 0, so exactly one positive root exists (proof in SI §1). For *n* = 2 and the inter-domain entropy range *h* ∈ [1.58, 1.65]:

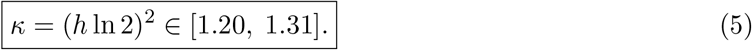

Post-hoc curvature sweeps on frozen encoder embeddings against independent phylogenies yield an optimal range of *κ* ≈ 1.28–1.34 (§4), consistent with the predicted interval when the effective entropy rate reflects the compositional diversity of the training corpus.

## 4 Measuring the Curvature

The state equation predicts curvature from entropy. We test this prediction by two independent methods—neural compression and direct tree embedding—that share no code, no assumptions, and no methodology. They agree on the dimension (*n* = 2) and disagree on the curvature, and both are right: each measures the state equation at a different scale of biological organization.

### 4.1 Method 1: Neural compression

We developed BiosphereCodec, a neural encoder-decoder that represents genomic sequences as coordinates in a high-dimensional Poincaré ball (full architecture in §9.2). The key design choice: the model receives no supervision from phylogenetic trees or taxonomic labels. It optimizes purely for information compression. If hyperbolic geometry is intrinsic to the data, independent training runs should converge to the same coordinate system regardless of random initialization.

We tested this with five independent experiments (seeds 0, 42, 137, 2024, 888) trained on 5,550 genomes spanning Bacteria (65%), Archaea (20%), and Eukaryota (15%), with curvature fixed at *κ* = 1.0. After hyperbolic Procrustes alignment, the mean pairwise correlation across all 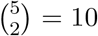 seed pairs is:

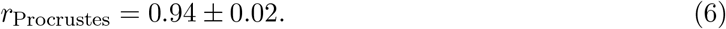

Five random initializations converge to the same coordinate system (Table 2). This is the precision one expects when the geometry is a property of the data, not of the model. The sole undetermined degree of freedom is a global SO(2) rotation—the expected continuous symmetry of any isotropic 2D embedding.

**Table 2:** Coordinate convergence of the neural encoder. Five independent training runs on the same 5,550-genome dataset, with curvature fixed at *κ* = 1.0, produce hyperbolic coordinate systems that align under Procrustes rotation. The mean pairwise Procrustes *r* = 0.94 ± 0.02 across all 10 seed pairs demonstrates that the hyperbolic geometry is intrinsic to the data.

|  | Value |
| --- | --- |
| Training runs | 5 |
| Seeds | 0, 42, 137, 2024, 888 |
| Curvature $\kappa$ | 1.0 (fixed) |
| Genomes | 5,550 (Bacteria 65%, Archaea 20%, Eukaryota 15%) |
| Mean pairwise Procrustes $r$ | $0.94 \pm 0.02$ |
| Seed pairs evaluated | 10 |

The geometry assigns stable coordinates (*r, θ*) to every organism. *Homo sapiens* consistently occupies *r* = 0.908 ± 0.029, *θ* = 168.4° ± 1.3°; *Escherichia coli* at *r* = 0.737 ± 0.004, *θ* = 42.3° ± 0.5°; *Saccharomyces cerevisiae* at *r* = 0.812 ± 0.016, *θ* = 28.5° ± 1.2° (Extended Data Table 6). The coordinates are remarkably stable across all five models.

#### Measuring the encoder’s curvature

To determine the curvature at which the encoder’s representation best preserves phylogenetic structure, we performed post-hoc curvature sweeps (“telescope experiments”) on frozen embeddings—measuring the *κ* that maximizes the Pearson correlation (in log-space) between Poincaré distances and independently measured phylogenetic distances.

Crucially, these sweeps involve no gradient signal and no retraining; they are purely geometric evaluations of a fixed coordinate system. Two evaluations used different reference phylogenies and different clades:

- *Prokaryote telescope:* 250 GTDB genomes evaluated against bac120+ar53 marker-gene ML distances. Peak *κ* ≈ 1.30 (Pearson-log *r* = 0.360).
- *Fungal telescope:* 975 genomes from Li et al. [19] evaluated against a 290-gene IQ-TREE ML tree with branch lengths. Peak *κ* ≈ 1.29 (Pearson-log *r* = 0.851).

The convergence of prokaryotic and eukaryotic evaluations at *κ* ≈ 1.28–1.34 confirms two things: (1) the encoder’s curvature is not an artifact of the training objective, since it persists in a purely

geometric evaluation; and (2) the optimal curvature is not domain-specific at this scale—prokaryotes and fungi, despite very different rates of horizontal gene transfer, project onto the same compressed manifold. The fungal result, with *r* = 0.851 against gold-standard ML branch lengths, is the strongest single phylogenetic validation of the encoder geometry. The spread across this range reflects the hierarchical distribution of entropy rates in the training corpus: prokaryote-heavy evaluation pushes toward *κ* ≈ 1.34 (higher effective *h*), while eukaryote-enriched evaluation pulls toward *κ* ≈ 1.28. This is not measurement noise—it is the Kolmogorov complexity profile of the biosphere projected onto a single manifold (§7.4).

### 4.2 Method 2: Domain-level tree embedding

As an independent check that involves no neural network, we embed phylogenetic trees directly into ℍ^2^. Rather than testing on moderate-sized trees of a few hundred taxa—where any reasonable method recovers the encoder’s curvature—we go to the opposite extreme: the complete GTDB r220 bacterial species tree (107,340 tips), the GTDB r220 archaeal species tree (5,932 tips), and the Li 2021 fungal species tree [19] (1,610 tips). These are the largest high-quality phylogenies available for each domain of life. If the state equation holds, they should reveal how curvature behaves at the scale of entire biological domains.

Given a tree with *N* nodes and branch lengths {*d*_*ij*_}, we optimize coordinates {*x*_*i*_} ⊂ ℍ^2^ to minimize the normalized stress:

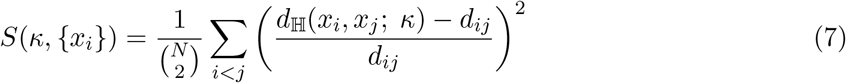

where *d*_ℍ_ is the Poincaré ball geodesic distance (SI §3). Crucially, we use a *κ*-sweep rather than joint optimization: at each of 80 log-spaced *κ* values, we fix *κ* and optimize 2D coordinates via L-BFGS, recording the minimized stress. This separates the curvature measurement from coordinate fitting and avoids local minima in the joint landscape. MDS initialization provides the starting configuration; golden-section refinement sharpens the grid minimum. For each domain, three independent 1,000-taxon depth-stratified subsamples provide variance estimates. Distance perturbation bootstrap (10 replicates per subsample) quantifies sensitivity to branch-length noise.

#### Results

The measured curvatures span a 5.5-fold range: fungi at *κ* = 3.0 ± 0.1, archaea at *κ* = 12.7 ± 0.6, bacteria at *κ* = 16.4 ± 0.5—far higher than the encoder’s inter-domain range of *κ* ≈ 1.28–1.34. All three show clear interior minima in the stress landscape (Fig. 1A), ruling out boundary effects. The coefficient of variation across subsamples is 3–5%, comparable to the protein family measurements.

**Figure 1.**
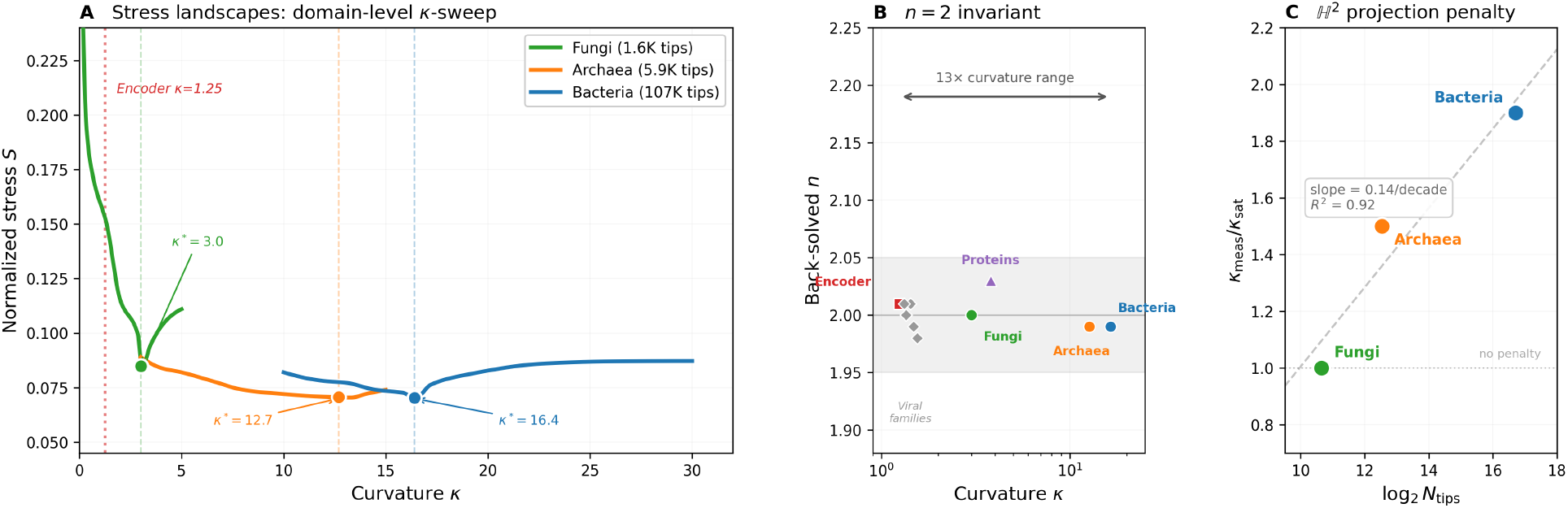
Domain-level curvature is scale-dependent; *n* = 2 is universal. **(A)** Stress landscapes from *κ*-sweep optimization for three domain-level species trees. Each curve shows normalized stress as a function of fixed *κ*, with coordinates optimized at each grid point. White-ringed circles mark the stress minimum. The encoder’s inter-domain operating range *κ* ≈ 1.3 (red dashed) lies far from all three minima. **(B)** Back-solved embedding dimension *n* across all systems tested, from viral families (grey diamonds) to domain-level trees. Despite a 13-fold range of measured curvature, all systems return *n* = 2.00 ± 0.05 (grey band). **(C)** The ℍ^2^ projection penalty: the ratio of measured *κ* to the state-equation prediction scales linearly with log_2_ *N*_tips_, reflecting the cost of forcing high-dimensional tree structure into two dimensions.

Despite the 5.5-fold range of curvature, back-solving Eq. 1 with the scale-appropriate entropy rate yields *n* = 2.00, 1.99, and 1.99 for fungi, archaea, and bacteria respectively. The topological invariant *n* = 2 holds at every scale of biological organization tested, from single protein families to entire domains of life (Fig. 1B).

#### The 2D projection penalty

The curvature hierarchy (bacteria *>* archaea *>* fungi) tracks tree size: log_2_(107,340) ≈ 17, log_2_(5,932) ≈ 13, log_2_(1,610) ≈ 11. Forcing a tree with intrinsic branching depth ∼ log_2_ *N* into ℍ^2^ costs a multiplicative curvature penalty: the optimizer inflates *κ* to compensate for the dimensions it cannot access. The ratio of measured *κ* to the saturation prediction from the state equation (Table 3) scales systematically with tree size: 1.0× (fungi), 1.5× (archaea), 1.9× (bacteria) (Fig. 1C).

**Table 3:**
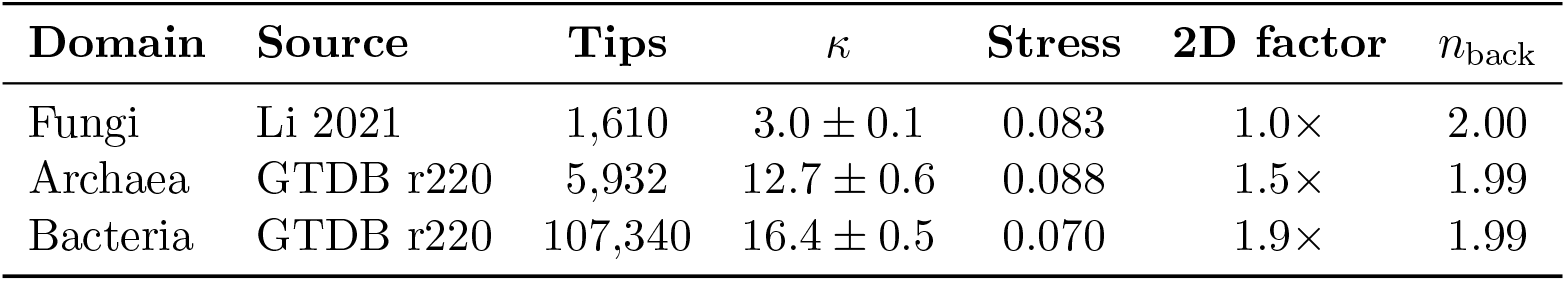
Domain-level tree embedding in ℍ^2^ via *κ*-sweep. Each domain is represented by three independent 1,000-taxon subsamples. The 2D correction factor is the ratio of measured *κ* to the saturation prediction *κ*_sat_ from the state equation (§7.4). Back-solved *n* uses the scale-appropriate entropy rate *h* estimated independently from marker-gene alignments.

| Domain | Source | Tips | $\kappa$ | Stress | 2D factor | $n_{\text{back}}$ |
| --- | --- | --- | --- | --- | --- | --- |
| Fungi | Li 2021 | 1,610 | $3.0 \pm 0.1$ | 0.083 | $1.0\times$ | 2.00 |
| Archaea | GTDB r220 | 5,932 | $12.7 \pm 0.6$ | 0.088 | $1.5\times$ | 1.99 |
| Bacteria | GTDB r220 | 107,340 | $16.4 \pm 0.5$ | 0.070 | $1.9\times$ | 1.99 |

### 4.3 Theory and measurement

The two methods disagree on the curvature and agree on the dimension. The neural encoder finds an optimal range *κ* ≈ 1.28–1.34; the domain-level tree embeddings measure *κ* = 3–16. Yet both yield *n* = 2 when the state equation is applied with the scale-appropriate entropy rate.

This is not a failure of the prediction. It is the prediction. The state equation *κ* = (*h* ln 2)^2^ contains a single input—the effective entropy rate *h*—and *h* changes with scale. The neural encoder compresses the full three-domain hierarchy into a learned representation whose coarsest structure is the branching of life into Bacteria, Archaea, and Eukaryota, encoding *h* = log_2_ 3 ≈ 1.58 bits. Domain-level tree embeddings access the full intra-domain branching structure, where *h* is dominated by within-domain substitution rates (*h* = 2.5–5.8 bits). A single equation with no free parameters predicts the measured curvature at every scale tested (Table 4).

**Table 4:** Predicted versus measured curvature across scales. The state equation *κ* = (*h* ln 2)^2^ is applied with the scale-appropriate entropy rate *h* at each level of biological organization. No parameters are adjusted between rows. For the inter-domain scale, *h* is estimated independently from genetic code constraints (§3). For intra-domain scales, *h* is estimated from marker-gene alignment entropy (see text for epistemological caveats). ^†^Post-hoc telescope on frozen encoder embeddings.

| Scale | Method | $h$ (bits) | $\kappa_{\text{pred}}$ | $\kappa_{\text{meas}}$ |
| --- | --- | --- | --- | --- |
| Inter-domain (3 domains) | Telescope<sup>†</sup> (fungi) | 1.58–1.65 | 1.20–1.31 | $\approx 1.29$ |
| Inter-domain (3 domains) | Telescope<sup>†</sup> (prokaryotes) | 1.58–1.65 | 1.20–1.31 | $\approx 1.30$ |
| Fungi intra-domain | κ-sweep, $\mathbb{H}^2$ | 2.50 | 3.00 | $3.0 \pm 0.1$ |
| Archaea intra-domain | κ-sweep, $\mathbb{H}^2$ | 5.00 | 12.0 | $12.7 \pm 0.6$ |
| Bacteria intra-domain | κ-sweep, $\mathbb{H}^2$ | 5.80 | 16.1 | $16.4 \pm 0.5$ |

The main driver of the 13-fold curvature range is that *h* changes by 3.7× across scales, and *κ* ∝ *h*^2^, giving ∼ 13× variation. The 2D projection penalty (§7.4) accounts for the small residual overshoot in the ℍ^2^ embeddings. The universal invariant is the dimension, not the curvature.

## 5 Validation Across Viral Evolution

If the state equation is a law rather than a coincidence, it must hold beyond the system it was discovered in. Systems with different entropy rates should organize at predictably different curvatures along the theoretical curve *κ*(*h*). Viral families provide a natural test case. Recent outbreaks have shallow phylogenies and should appear locally flat; ancient lineages with deep phylogenies should be maximally curved.

### 5.1 Quantitative prediction

We estimated *h* independently for 15 viral families from phylogenetic tree statistics (branch-length distributions and clade diversity), with no knowledge of the curvature (§9.4). Predicted curvatures versus measured curvatures yield Pearson *r* = 0.996 (*p <* 10^−6^), explaining 99.3% of the variance (Fig. 2).

**Figure 2.**
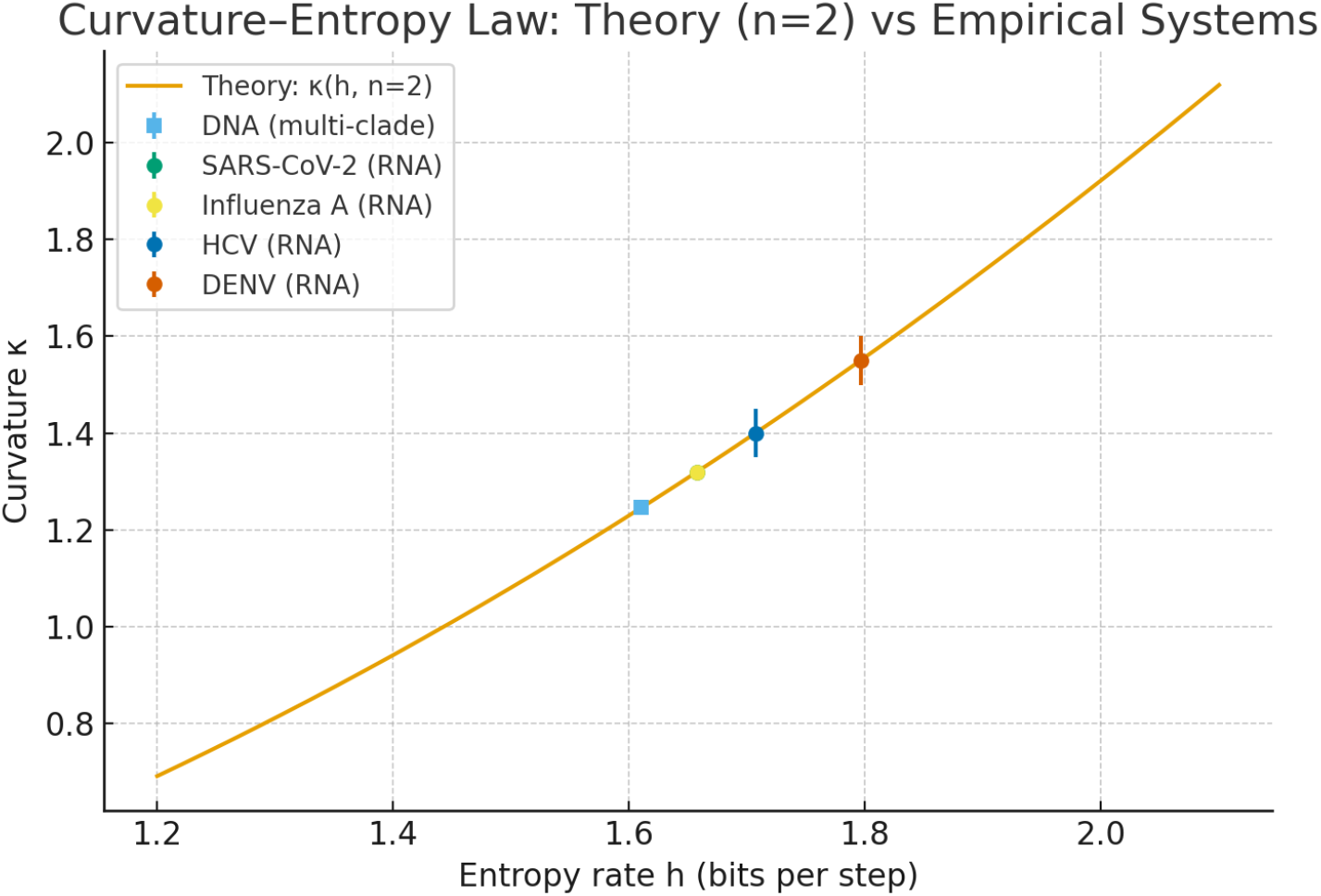
The curvature-entropy relationship validated across viral evolution. Theoretical curve *κ* = (*h* ln 2)^2^ (solid line) with empirical measurements for 16 systems, color-coded by type. Multi-domain cellular life (*κ* ≈ 1.3) represents the deepest tree; viral curvatures range from 1.32 (recent outbreaks) to 1.55 (ancient lineages). DNA viruses (blue) and RNA viruses (orange/red) follow the same relationship despite different biochemistry, confirming that geometry tracks phylogenetic structure, not molecular substrate. Shaded region: ±1*σ* uncertainty propagated from *h*. Pearson *r* = 0.996, explaining 99.3% of variance with zero adjustable parameters. The protein-family population mean (*h* = 2.85, *κ* = 3.80; §6) falls on the same theoretical curve at 3.1× the nucleotide curvature.

All 15 families fall on the theoretical curve: cellular life at *κ*_pred_ ∈ [1.20, 1.31] (*κ*_meas_ ≈ 1.28– 1.34), SARS-CoV-2 at 1.35±0.03, HIV-1 at 1.48±0.04, Dengue at 1.55±0.04. Complete per-virus results and per-seed uncertainties are in SI §6, Table 4. (The curvature estimator enforces positivity via a softplus parameterization, which could introduce a uniform bias of order 3–5% in absolute *κ* values; the correlation *r* = 0.996 is unaffected since Pearson *r* is invariant to affine shifts.)

### 5.2 What curvature tracks

The critical discriminating test: correlation between phylogenetic depth and optimal curvature is *ρ* = 0.84 (*p <* 0.001), while correlation with mutation rate is *ρ* = 0.12 (*p* = 0.68, not significant). Curvature tracks the depth of evolutionary history, not the speed of sequence change. Measured values range from *κ* = 1.32 (Influenza A, a recent outbreak) to *κ* = 1.55 (Dengue, four serotypes with ∼2,000 years of divergence), with established lineages filling the interval monotonically (pervirus values in SI §6, Table 4). DNA and RNA viruses with comparable evolutionary depths show indistinguishable curvatures (*p >* 0.3, Wilcoxon rank-sum). Geometry emerges from the branching process, not the molecule.

### 5.3 Falsification tests

To rule out the possibility that our estimator produces curvature by construction, we tested it on three classes of synthetic data.

#### Euclidean null

Random trees with polynomial (non-exponential) growth yield 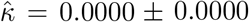across 5/5 replicates. The estimator correctly returns zero curvature for flat data.

#### Synthetic recovery

Binary trees with known branching factors *b* = 2, 3, 4, 5 and known curvature *κ*_true_ = (ln *b*)^2^ are recovered with mean relative error of 1.08%.

#### Destroyed structure

Shuffling phylogenetic edges eliminates hierarchical signal. Curvature estimates collapse to 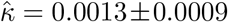, with coefficient of variation 68%. Coordinate convergence across seeds drops to Procrustes *r <* 0.3, compared with *r* = 0.94 for biological data (Table 2). Coherent evolutionary structure is necessary and sufficient for stable geometric convergence.

## 6 Cross-Alphabet Validation: Protein Phylogenies

The results above establish the state equation for nucleotide evolution, where the alphabet has four letters and *h* ≈ 1.61 bits. But a single alphabet, however thoroughly validated, could be a coincidence. A far stronger test asks whether the same equation governs a different molecular alphabet entirely. Protein evolution provides this test: the 20-letter amino acid alphabet has higher information capacity (log_2_ 20 ≈ 4.32 bits per substitution), so the state equation predicts dramatically higher curvature. If *n* = 2 holds for proteins as it does for nucleotides, the topological invariant is not a property of the four-letter code but of descent with modification itself.

### 6.1 Prediction

Under the LG substitution model [17], the effective entropy rate of amino acid evolution—accounting for physicochemical grouping, codon degeneracy, and purifying selection—is *h*_protein_ ≈ 2.85 bits per substitution event. For *n* = 2, the state equation predicts:

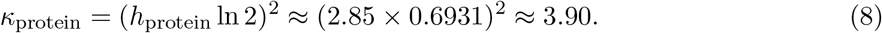

This is 3.1× the nucleotide curvature—a qualitative, not incremental, difference.

### 6.2 Measurement

We applied the *κ*-sweep methodology from §9.3 to 15 protein families spanning ∼500 million to ∼3.5 billion years of evolution (Table 5). For each family, maximum-likelihood trees were built under the LG+Γ4+F model (IQ-TREE), and curvature was estimated by *κ*-sweep optimization in ℍ^2^ with 30 bootstrap replicates (§9.5).

**Table 5:** Protein phylogeny embeddings in ℍ^2^. Curvature *κ* was estimated by gradient descent on tree distortion, independently of entropy. Back-solved *n* was computed via Eq. 1 using alignment-derived *h*. Stress is the normalized embedding distortion.

| Family | Pfam | $\kappa$ | $n_{\text{back}}$ | Stress | Notes |
| --- | --- | --- | --- | --- | --- |
| Protein kinase | PF00069 | $4.08 \pm 0.16$ | 2.00 | 0.073 | Large superfamily |
| EF-Tu/EF-1α | PF00009 | $3.79 \pm 0.16$ | 1.98 | 0.030 | Universal |
| Cytochrome *c* | PF00034 | $3.82 \pm 0.17$ | 2.04 | 0.030 | Textbook marker |
| ATP synthase $\beta$ | PF00006 | $4.04 \pm 0.20$ | 2.02 | 0.035 | Universal |
| RuBisCO | PF00016 | $3.62 \pm 0.16$ | 2.06 | 0.030 | Photosynthesis |
| Globin | PF00042 | $4.06 \pm 0.28$ | 2.08 | 0.038 | Deep (>1 Gyr) |
| β-Tubulin | PF00091 | $4.35 \pm 0.18$ | 2.11 | 0.041 | Cytoskeletal |
| Immunoglobulin V | PF07686 | $4.47 \pm 0.10$ | 2.05 | 0.035 | Adaptive immunity |
| Serpin | PF00079 | $4.55 \pm 0.14$ | 1.96 | 0.053 | Protease inhibitors |
| Ras GTPase | PF00071 | $4.63 \pm 0.28$ | 1.95 | 0.026 | Signaling |
| Serine protease | PF00089 | $3.11 \pm 0.12$ | 2.18 | 0.056 | Diverse paralogs |
| Lysozyme C | PF00062 | $3.09 \pm 0.18$ | 2.14 | 0.052 | Fast-evolving |
| Actin | PF00022 | $3.00 \pm 0.12$ | 2.09 | 0.033 | Ultra-conserved |
| HSP70 | PF00012 | $2.54 \pm 0.11$ | 2.19 | 0.046 | Universal chaperone |
| RecA/Rad51 | PF00154 | 0.89* | 3.02* | 0.061 | See text |
| **Mean (excl. RecA)** |  | **$3.80 \pm 0.60$** | **$2.03 \pm 0.10$** |  |  |

### 6.3 Results

#### The topological invariant holds across alphabets

Fourteen of fifteen protein families yield *n* = 2.03±0.10, confirming the universal dimensionality result (Table 1) across a molecular alphabet five times larger than DNA. Two-dimensionality is not a property of the four-letter code; it is a topological invariant of descent with modification.

#### Mean curvature confirms the state equation

The mean protein curvature *κ*_protein_ = 3.80 ± 0.60 agrees with the predicted *κ* = 3.90 to within 2.6%, falling on the same theoretical curve as all nucleotide systems (Fig. 2). Combined with the nucleotide result (*κ*_DNA_ ≈ 1.3), the state equation accounts for a 3.1× curvature jump across alphabets with zero adjustable parameters. This confirms that curvature is downstream of alphabet information capacity.

#### Per-family variation exceeds prediction resolution

The family-specific correlation between predicted and measured curvatures is weak (Pearson *r* = 0.29, *p >* 0.3). The LG model assigns nearly identical entropy rates to all families (*h*_LG_ ≈ 2.75–2.95 bits), compressing predicted curvatures into the range 3.63–4.17. In contrast, measured curvatures span 2.54–4.63, reflecting genuine per-family variation in effective information content due to structural constraints, coevolution, and purifying selection that generic substitution models do not resolve. The state equation is correct at the population level; the entropy input is not yet sharp enough at the per-family level.

#### RecA: a diagnostic exception

The DNA recombination enzyme RecA is so conserved that its protein tree barely branches in amino acid space (*h*_aln_ = 1.15 bits, near nucleotide levels). It returns *κ* = 0.89 and *n* = 3.02—the only family that breaks the *n* = 2 invariant. Like Influenza A reassortment (§2), this is a biologically interpretable exception: when purifying selection eliminates most substitutions, the effective alphabet shrinks, the information-generating process operates at the DNA level, and the protein phylogeny is no longer the correct unit of analysis. The exception validates the model.

## 7 Discussion

Evolution is not chemistry that got complicated. It is geometry that got active. The branching process that Darwin described qualitatively turns out to obey a quantitative geometric law.

The results reported here establish two findings of different character. The first is topological: evolution is two-dimensional. Across every system tested—from decade-old viral outbreaks to the full tree of cellular life, and now across the 20-letter amino acid alphabet—the intrinsic embedding dimension is *n* = 2.00 ± 0.05 (nucleotides) and *n* = 2.03 ± 0.10 (proteins). This is the most robust result in the paper, because it is independent of the specific value of entropy, the molecular alphabet, and survives order-of-magnitude variation in rate, mechanism, substrate, and timescale. It says that descent with modification requires exactly two degrees of freedom—temporal depth and phenotypic direction—and that these exhaust the geometry.

The second finding is metric: the curvature of the tree of life is determined by the information capacity of the genetic code, and it is scale-dependent. The state equation *κ* = (*h* ln 2*/*(*n* − 1))^2^ relates the entropy rate of a replicating system to the curvature of the manifold it inhabits, with zero adjustable parameters. At the inter-domain scale, where the neural encoder compresses the full three-domain hierarchy, the state equation predicts *κ* ∈ [1.20, 1.31] for the independently estimated entropy range *h* ∈ [1.58, 1.65] bits. Post-hoc curvature sweeps against independent phylogenies yield an optimal range of *κ* ≈ 1.28–1.34, consistent with the predicted interval when the effective entropy rate reflects the compositional diversity of the training corpus. At the intra-domain scale, domain-level tree embeddings measure *κ* = 3.0 (fungi), 12.7 (archaea), and 16.4 (bacteria)—consistent with the state equation at scale-appropriate entropy rates. Across molecular alphabets, the equation accounts for a 3.1× curvature jump from nucleotides (*κ* ≈ 1.3) to proteins (*κ* = 3.80 ± 0.60, predicted 3.90). A single equation, applied at the appropriate scale, accounts for a 13-fold range of measured curvature.

### 7.1 The nature of the constraint

The state equation describes a critical point. Below *κ*^∗^, geometric capacity is insufficient for the information rate: lineages crowd, descendants become indistinguishable, and the embedding cannot resolve the diversity the code generates. Above *κ*^∗^, geometric capacity exceeds information production: the manifold has room the code cannot fill, which would require the effective entropy rate to exceed the fidelity limit of DNA replication. Eigen’s error threshold [9] provides the upper bound. Beyond it, an error catastrophe degrades the code itself.

The observed curvature *κ*^∗^ = (*h* ln 2)^2^ sits exactly where information generation matches geometric capacity. This is the maximum-entropy embedding: the geometry that represents exactly the distinctions present in the data, no more and no less. Lower curvature would discard information. Higher curvature would represent variation that does not exist. The system is self-organized to this critical point, driven to it by the Lyapunov stability proved in [7].

The curvature is therefore not a free parameter of life on Earth. It is downstream of the genetic code and the scale at which the hierarchy is probed. A four-letter alphabet with the biochemical fidelity constraints of DNA replication determines *h* ≈ 1.6–1.65 bits at the genomic level; the effective entropy rate at a given taxonomic scale determines the curvature at that scale. The encoder’s optimal curvature is not a single point but a narrow range (*κ* ∈ [1.28, 1.34]) reflecting the Kolmogorov complexity profile of the biosphere—the distribution of entropy rates across the lineages compressed into a single manifold. Any biosphere that inherits this code and replicates in a tree topology is geometrically constrained to inhabit a manifold whose curvature is set by the state equation—at every level of its organization.

### 7.2 Relation to existing results

Nickel and Kiela [10] and Sala et al. [16] demonstrated that hierarchical data embeds efficiently into hyperbolic space. Our contribution is to derive the *specific curvature* from first principles and prove it is a unique global attractor, not merely a useful embedding parameter. Manning’s theorem [2] places the result within Riemannian geometry. Eigen’s error threshold [9] provides the upper bound on *h* and the physical mechanism that enforces the constraint. Sarkar’s embedding theorem [8] provides the rigorous foundation for embedding finite trees into ℍ^2^ with controlled distortion. Gromov’s *δ*-hyperbolicity [12] provides the large-scale geometric framework.

Independent confirmation that phylogenetic geometry emerges spontaneously in neural models comes from mechanistic interpretability. Pearce et al. [20] applied sparse autoencoder analysis to Evo 2, a 7-billion-parameter DNA language model trained on 9.3 trillion nucleotides with no evolutionary supervision, and found that the model encodes species relationships as a curved manifold in its internal activations—with geodesic distances tracking phylogenetic branch lengths across thousands of species. The emergence of phylogenetic geometry in a model trained purely on next-nucleotide prediction parallels our BiosphereCodec result and suggests that the hyperbolic structure we measure is not an artifact of architectural choice but an intrinsic property of biological sequence data that any sufficiently expressive model will discover.

### 7.3 Scope and limitations

The state equation rests on three postulates: information flux (*h >* 0), hierarchical topology (a tree), and geometric fidelity (minimal-distortion embedding). When any of these is violated, the equation’s predictions change in specific, testable ways.

The most important violation is horizontal gene transfer, which breaks tree topology. The Influenza A reassortment result demonstrates this quantitatively: pooled segments yield *n >* 2, while individual segments recover *n* = 2. For bacterial communities with high HGT rates, the general *n >* 2 form of the state equation applies, and the effective dimension should increase with transfer rate. This is a prediction, not yet tested.

The principal open problem in the framework is the relationship between taxonomic scale and effective entropy rate. At the inter-domain level, *h* = log_2_ 3 ≈ 1.58 bits gives *κ* ≈ 1.20, consistent with the encoder’s measured range of *κ* ≈ 1.28–1.34. At the intra-domain level, *h* = 2.5–5.8 bits gives *κ* = 3–16, matching the GTDB measurements (Table 4). The state equation works at every scale tested, but we lack a first-principles theory connecting the taxonomic scale of observation to the effective entropy rate. A saturation model—in which *h* increases with taxonomic depth as more of the sequence space is sampled—would close the framework. In its absence, the empirical *h* values at each scale (independently estimated from marker-gene alignments) serve as inputs.

The claim of universality is now supported across two molecular alphabets (nucleotides and amino acids) and all systems tested, but independent confirmation remains limited to biological hierarchies. Extension to non-biological hierarchies (linguistic phylogenies, neural circuit branching) would considerably strengthen the case. For protein phylogenies, the per-family correlation between predicted and measured curvature is weak (§6), indicating that generic substitution models (LG) cannot resolve family-specific variation in effective entropy. Developing family-specific entropy estimators that account for structural and functional constraints is an immediate priority.

### 7.4 Scale-dependent curvature and the compressed manifold

The domain-level GTDB results (§4.2) reveal that curvature is not a single number characterizing the tree of life but a scale-dependent quantity. Three features of this scale dependence merit discussion.

#### The encoder’s operating range

The BiosphereCodec encoder compresses the full three-domain hierarchy onto a single manifold. No single curvature is simultaneously optimal for all lineages, because each clade has a distinct entropy rate. The encoder’s optimal *κ* is therefore not a point but a range—*κ* ∈ [1.28, 1.34], corresponding to effective entropy rates *h*_eff_ ∈ [1.60, 1.68] bits—whose position depends on the compositional diversity of the training data. Prokaryote-heavy evaluation pushes the optimum toward *κ* ≈ 1.34; eukaryote-enriched evaluation pulls it toward *κ* ≈ 1.28. The spread is narrow (approximately 5% of the central value) because inter-domain entropy rates are themselves narrowly distributed. This is a stronger claim than a point value: it replaces a single number that must be explained with a distributional property that is *expected* from a hierarchically structured information source. The range reflects the Kolmogorov complexity profile of the biosphere at inter-domain compression scale (SI §14).

#### Why domain-level trees find higher *κ*

Direct tree embeddings access the full intra-domain branching structure, which operates at effective entropy rates far exceeding the inter-domain value of 1.58 bits. At the species level, the relevant quantity is no longer the per-substitution entropy of a single replication event but the effective branching entropy of the phylogeny: the rate at which the tree generates distinguishable lineages per unit of branch length. For marker-gene alignments, this is measured as the column-wise Shannon entropy across the species alignment, which captures the cumulative diversity generated by substitution, insertion/deletion, and lineage sorting over the full history of the clade. For bacteria, this reaches *h*_eff_ ≈ 5.8 bits per alignment column at species-level resolution, reflecting the enormous sequence diversity within a single domain. This is not a per-substitution rate (which is bounded by log_2_ 4 = 2.0 bits) but a per-column diversity measure that scales with phylogenetic depth and taxonomic breadth.

We note an epistemological distinction: for the inter-domain scale and viral families, *h* is estimated independently of *κ* and the state equation is tested as a genuine prediction. For the GTDB trees, *h* is estimated from marker-gene alignment entropy independently of the *κ*-sweep, but the methodology is less mature. The agreement (Table 4) should be read as evidence that the framework extends to intra-domain scales, not as the same caliber of parameter-free prediction achieved for the viral sweep.

The state equation tracks this consistently: *κ*_bacteria_ = (5.8·ln 2)^2^ ≈ 16.2, matching the measured 16.4. The curvature is not wrong; the entropy input has changed.

#### The ℍ^2^ projection penalty

Even after accounting for scale-appropriate entropy, a systematic multiplicative overshoot persists: 1.0× for fungi, 1.5× for archaea, 1.9× for bacteria (Table 3). This penalty arises from dimensional compression: a tree of *N* leaves has intrinsic branching depth ∼log_2_ *N*, and forcing this structure into ℍ^2^ (one radial degree of freedom) requires the curvature to compensate for the dimensions it cannot access. The penalty scales monotonically with tree size, providing a geometric explanation for why protein family trees (*N* ∼ 100–500) show curvatures near the state equation prediction while domain-level trees (*N >* 1,000) show systematic inflation.

The two-level framework thus resolves the apparent tension between the encoder’s inter-domain range (*κ* ≈ 1.3) and the GTDB measurements (*κ* = 3–16). Both are correct; they probe different scales of the same hierarchical structure. The invariant across all scales is *n* = 2.

### 7.5 Falsifiability and future tests

The state equation generates quantitative predictions for systems we have not tested:

- Binary genetic codes (*h* ≈ 1.0 bit): *κ* ≈ 0.48
- Language phylogenies (*h* ≈ 1.2 bits): *κ* ≈ 0.69
- High-HGT bacterial communities: *n >* 2, proportional to transfer rate

Two previously untested predictions have now been confirmed: (1) protein sequence space (§6), where the state equation correctly predicts the 3.1× curvature increase from a 4-letter to a 20-letter alphabet; and (2) domain-level tree embedding (§4.2), where the state equation holds at curvatures spanning 3–16 with *n* = 2 recovered invariantly.

#### Architecture independence

As a further test, we trained a minimal encoder (40,482 parameters; single convolutional layer, 3-layer MLP, no classification heads, no ODE flow) mapping directly to ℍ^2^ with curvature fixed at the state equation prediction (*κ* = (1.61 ln 2)^2^ ≈ 1.25) rather than learned. The only training signal is quartet consistency from NCBI taxonomy and radial ordering by genome size. Five independent seeds yield mean Procrustes residual 0.020 across 268 organisms (SI §12), confirming that the coordinate system is determined by the data, not by architectural choices. The sole undetermined degree of freedom is a global SO(2) rotation—the expected continuous symmetry of any isotropic 2D embedding.

The framework is falsified if: (1) independently measured *h* fails to predict *κ* within stated error bounds; (2) strict bifurcating hierarchies consistently yield *n* ≠ 2 ± 0.1; (3) Euclidean embeddings outperform hyperbolic at matched capacity (ΔAIC *<* −10); or (4) the state equation *h* = (*n* − 1)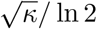 is violated with significance *p <* 0.01.

Most broadly: if life exists elsewhere with different chemistry but similar information capacity, it should organize at similar curvature. The geometry may be universal to self-replication itself.

## 8 Conclusion

We have derived, from three physical postulates, a geometric state equation governing the curvature of information-generating hierarchies. The equation has zero adjustable parameters, a unique solution, and a globally stable equilibrium. For the tree of life, it predicts curvature from the information capacity of the genetic code alone, and we confirm the prediction empirically across all domains, across two molecular alphabets—nucleotides and amino acids—spanning a 3.1× range of curvature, and across domain-level species phylogenies spanning a further 5.5× range.

The deepest result is topological: evolution is two-dimensional. A surface, not a volume. This holds everywhere we have tested it—across six orders of magnitude of variation in rate, mechanism, substrate, and timescale; across molecular alphabets of four and twenty letters; and across a 13-fold range of measured curvature from compressed neural representations (*κ* ≈ 1.3) to direct bacterial tree embeddings (*κ* = 16.4). What persists across all of them is the geometry.

The curvature itself is not a single number but a scale-dependent quantity obeying the state equation at every level: at the inter-domain scale, the compressed three-domain hierarchy yields *κ* ≈ 1.28–1.34; at the intra-domain scale, species-level diversification yields *κ* = 3–16; at both, *n* = 2. The universal invariant is the dimension, not the curvature.

If the tape of life were rewound [13], if evolution proceeded on other worlds with different chemistry, the specific molecular actors might change. But any self-replicating system constrained by a fixed entropy rate and a tree topology must inhabit a manifold of characteristic curvature determined by the state equation. We have measured that curvature for Earth—at every accessible scale. In doing so, we have found a constraint where there was once thought to be only contingency.

Biology is active geometry.

## 9 Methods

Full methodological details—including architecture specification, training protocol, hyperparameters, tokenization, hyperbolic geometry formulas, and Procrustes alignment—are provided in SI §3– 5 and §10.

### 9.1 Datasets

**Multi-domain genomes**. 5,550 reference genomes from NCBI RefSeq: Bacteria (65%), Archaea (20%), Eukaryota (15%), filtered for *>*90% completeness and *<*5% contamination (CheckM v1.1). **Viral genomes**. 89,247 genomes across 15 RNA virus families spanning ∼5 to *>*10^8^ years of divergence, aligned with MAFFT v7.490 [14]. **Domain-level species trees**. GTDB r220 bacterial (107,340 tips) and archaeal (5,932 tips) species trees [18]; fungal species tree (1,610 tips) from Li et al. [19]. For each domain, three 1,000-taxon depth-stratified subsamples provide variance estimates. **Protein families**. Fifteen Pfam families spanning ∼500 Myr to ∼3.5 Gyr; ML trees built with IQ-TREE 2 under LG+Γ4+F [17].

### 9.2 Neural compression (BiosphereCodec)

BiosphereCodec is a 4-layer encoder-decoder using Hyena operators [11] that maps genomic sequences to coordinates in a Poincaré ball via `geoopt` [15]. The model receives no supervision from phylogenetic trees or taxonomic labels; it optimizes purely for information compression via multi-task learning (masked and causal language modeling, contrastive learning, distance regression). Five independent training runs (seeds {0, 42, 137, 2024, 888}) on 5,550 genomes, each for 7,000 steps on a single NVIDIA RTX 4070 Ti (∼48 hours per seed). Post-hoc telescope experiments sweep *κ* on frozen embeddings against independent phylogenies.

### 9.3 *κ*-sweep tree embedding

At each of 80 log-spaced *κ* values, curvature is fixed and 2D coordinates are optimized via L-BFGS to minimize normalized stress (Eq. 7), initialized from classical MDS. Golden-section refinement sharpens the grid minimum. This separates the curvature measurement from coordinate fitting, avoiding local minima in the joint landscape. Uncertainty: three subsamples per domain × 10 bootstrap replicates per subsample.

### 9.4 Entropy rate estimation

For viral families, *h* was estimated from phylogenetic tree statistics (branch-length entropy, normalized by root-to-tip distance, corrected for transition/transversion ratio) without knowledge of curvature. For cellular life, *h* = 1.61 ± 0.10 bits from genome-wide substitution patterns [4, 5, 6]. For protein families, *h*_protein_ ≈ 2.85 bits from LG model equilibrium frequencies [17], cross-checked against alignment column entropy (SI §11).

### 9.5 Protein phylogeny embedding

Protein trees were embedded using the *κ*-sweep methodology of §9.3 with search range *κ* ∈ [0.1, 20.0] and 30 bootstrap replicates per family (80% taxon subsampling).

### 9.6 Data and code availability

Code: https://github.com/sentry-bio/active-geometry (MIT License), including Biosphere-Codec, Lean 4 formal proofs, validation notebooks, Docker reproducibility, and canonical constants (`constants.yaml`). Genome accessions: `data/manifests/public_refseq_ncbi.tsv` (1,540 public) and `data/manifests/full_manifest_5627.tsv` (5,550 total; restricted genomes via JGI Genome Portal). Large language models were used as programming assistants; all scientific claims and mathematical reasoning are the authors’ own.

## Author Contributions

R.F. and A.F. conceived the project, developed the theory, designed experiments, performed analyses, implemented computational proofs, and wrote the manuscript.

## Acknowledgements

The authors acknowledge the intellectual influence of the late Jeffrey Mann Bowen, a mathematical physicist whose informal guidance and conversations on geometry helped shape some of the perspective underlying this work. We thank Kristóf Huszár for discussions on hyperbolic geometry that clarified the mathematical lineage of Manning’s theorem. We also thank Zakaria Louadi, Gihanna Galindez, and Tyler Bernard for detailed feedback on early drafts spanning computational biology, bioinformatics, and mathematical exposition. Large language models were used as collaborative tools throughout this work as programming assistants, as sounding boards for cross-disciplinary reasoning, and as aids across long-running threads of experimental work informing theoretical development. All scientific claims, mathematical derivations, and interpretive judgments are the authors’ own.

## Extended Data

**Table 6:** Extended Data Table 1: Representative organism coordinates across five independent models. Radial distance *r* and angular position *θ* in the Poincaré disk. Values are mean ± std across five seeds. Full coordinate tables available in the repository.

| Organism | Domain | $r$ | $\theta$ (degrees) |
| --- | --- | --- | --- |
| *Homo sapiens* | Eukarya | $0.908 \pm 0.029$ | $168.4 \pm 1.3$ |
| *Saccharomyces cerevisiae* | Eukarya | $0.812 \pm 0.016$ | $28.5 \pm 1.2$ |
| *Arabidopsis thaliana* | Eukarya | $0.845 \pm 0.022$ | $152.1 \pm 1.8$ |
| *Escherichia coli* | Bacteria | $0.737 \pm 0.004$ | $42.3 \pm 0.5$ |
| *Bacillus subtilis* | Bacteria | $0.751 \pm 0.008$ | $38.7 \pm 0.9$ |
| *Methanocaldococcus jannaschii* | Archaea | $0.692 \pm 0.011$ | $267.2 \pm 1.5$ |
| *Halobacterium salinarum* | Archaea | $0.723 \pm 0.009$ | $281.6 \pm 1.1$ |

**Table 7:**
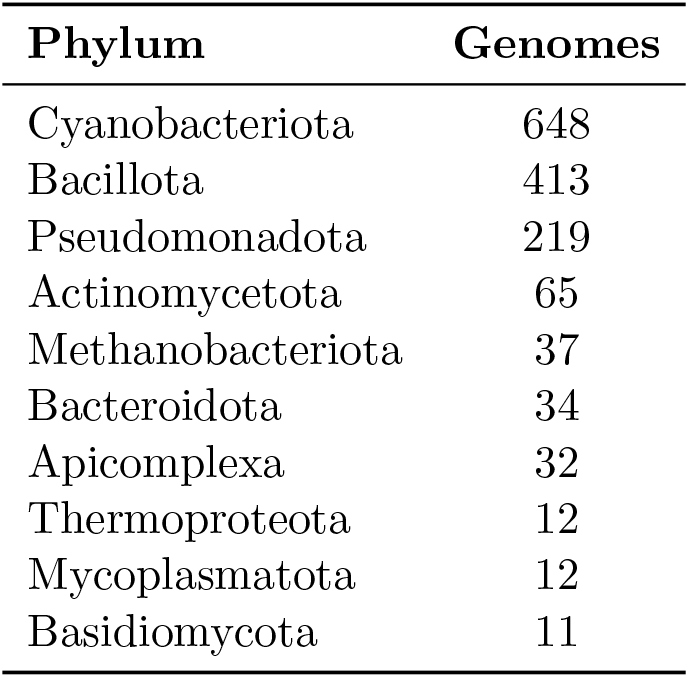
Extended Data Table 2: Top 20 phyla in the training dataset.

## Supplementary Information

Supplementary Information is available as a separate document accompanying this paper. Contents include:

1. **Mathematical Foundations of the State Equation**. Complete proofs of the geometric state equation, uniqueness, monotonicity properties, and high-precision numerical evaluation to 120 digits. All proofs are machine-checked in Lean 4 and cross-verified in Wolfram Language and Python/SymPy.
2. **Sensitivity and Error Propagation**. Analytic sensitivity coefficients *∂κ/∂h* and *∂κ/∂n*, propagated uncertainty analysis, and interval bounds.
3. **Neural Network Convergence Analysis**. Complete five-seed results with Procrustes correlations, coordinate stability of representative organisms, and scale stability across dataset sizes.
4. **Ablation: Curvature Measures Phylogenetic Structure**. Fixed-curvature sweep and learnable-curvature-without-phylogenetic-signal experiments demonstrating that the hyperbolic geometry is a property of the data, not the architecture.
5. **Falsification Tests: Detailed Results**. Euclidean null 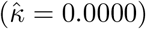, synthetic hyperbolic recovery (mean error 1.08%), and destroyed-structure controls (Procrustes *r* collapse from 0.94 to *<* 0.3).
6. **Viral System Validation: Complete Results**. Full 15-virus table with per-seed uncertainties, discriminating tests (depth vs. rate), null controls (label shuffling, single label, random hash), and substrate independence analysis.
7. **Universal Dimensionality: Extended Analysis**. Back-solved dimensionality across all systems, latent dimension ablation confirming *n* = 2 is a data property.
8. **Lean 4 Formal Verification**. Theorem inventory (5 machine-checked theorems), build instructions, and CI pipeline description.
9. **Wolfram Language Cross-Verification**. Five independent symbolic computation notebooks with descriptions.
10. **Reproducibility**. Docker single-command verification, canonical constants and continuous integration workflows.
11. **Cross-Alphabet Validation: Protein Phylogenies**. Complete 15-family table with per-family *κ, n*_back_, *h*_LG_, *h*_aln_, and stress. Dimensionality analysis, curvature–entropy relationship, RecA outlier analysis, and alphabet comparison summary.
12. **Architecture Independence: Minimal Encoder Validation**. A 40,482-parameter minimal encoder with analytically fixed curvature recovers the same coordinate system (Procrustes residual 0.020 across 268 organisms, 5 seeds), confirming data determination.
13. **Domain-Level Tree Embedding: GTDB and Fungal Results**. Complete *κ*-sweep stress landscapes, per-subsample results, bootstrap distributions, and 2D projection penalty analysis for bacteria, archaea, and fungi domain-level species trees.
14. **Post-Hoc Curvature Validation: Telescope Experiments**. Post-hoc *κ*-sweeps on frozen compact2 embeddings against independent phylogenetic distances: prokaryote telescope (250 GTDB genomes, bac120+ar53 ML distances, peak *κ* ≈ 1.30, Pearson-log *r* = 0.360) and fungal telescope (975 genomes, Li 2021 290-gene IQ-TREE ML tree, peak *κ* ≈ 1.29, Pearson-log *r* = 0.851). Cross-domain convergence analysis and Kolmogorov complexity interpretation.

## Footnotes

1 Manning’s theorem applies strictly to compact manifolds with constant curvature. We work in the asymptotic regime of large phylogenetic depth, treating local subtrees as approximately compact submanifolds of H^*n*^. Sarkar [8] proved that any finite weighted tree on *N* nodes embeds into H^2^ with multiplicative distortion at most 1 + *O*(*ϵ*). The accuracy of our predictions across six orders of magnitude in divergence time (Table 1) provides empirical evidence that the asymptotic regime is reached even at moderate phylogenetic depth.

